# The Bias of Using Cross-Validation in Genomic Predictions and Its Correction

**DOI:** 10.1101/2023.10.03.560782

**Authors:** Yanzhao Qian, Dinghao Wang, Qi Xuan Ding, Matthew Greenberg, Quan Long

## Abstract

Cross-validation (CV) is a widely used technique in statistical learning for model evaluation and selection. Meanwhile, various of statistical learning methods, such as Generalized Least Square (GLS), Linear Mixed-Effects Models (LMM), and regularization methods are commonly used in genomic predictions, a field that utilizes DNA polymorphisms to predict phenotypic traits. However, due to high dimensionality, relatively small sample sizes, and data sparsity in genomic data, CV in these scenarios may lead to an underestimation of the generalization error. In this work, we analyzed the bias of CV in eight methods: Ordinary Least Square (OLS), GLS, LMM, Lasso, Ridge, elastic-net (ENET), and two hybrid methods: one combining GLS with Ridge regularization (GLS+Ridge), and the other combining LMM with Ridge regularization (LMM+Ridge). Leveraging genomics data from the 1,000 Genomes Project and simulated phenotypes, our investigation revealed the presence of bias in all these methods. To address this bias, we adapted a variance-structure method known as Cross-Validation Correction (CVc). This approach aims to rectify the cross-validation error by providing a more accurate estimate of the generalization error. To quantify the performance of our adapted CVc towards all these methods, we applied the trained model to an independently generated dataset, which served as a gold standard for validating the models and calculating the generalization error. The outcomes show that, by leveraging CVc, we corrected the CV bias for most of the methods mentioned above, with two exceptions that are unrectifiable methods: ENET and Lasso. Our work revealed the substantial bias in the use of CV in genomics, a phenomenon under-appreciated by the field of statistical genomics and medicine. Additionally, we demonstrated that bias-corrected models may be formed by adapting CVc, although more work is needed to cover the full spectrum.

## 1 Introduction

Supported by modern high-throughput instruments, numerous genomic data have been collected and analyzed, leading to significant advancements in our understanding of inheritable traits. To date, over 5,700 Genome-Wide Association Studies (GWAS) have been undertaken, encompassing more than 3,300 traits and involving over a million participants. This extensive research has unveiled numerous associated and reproducible genetic variants that influence a range of inheritable traits (Watanabe et al., 2019). Concurrently, the availability of genomic data has facilitated the efforts of forming DNA-based phenotype predictors using various statistical models, such as Linear Mixed-Effects Models (LMM) (Yang et al., 2010; Zhang et al., 2010), regularization Methods (Zeng, Zhou, & Huang, 2017) and Bayesian methods (Zhou, Carbonetto, & Stephens, 2013). (In the context of genomic prediction, the estimation of random effects for LMM is referred to as gBLUP (Clark & van der Werf, 2013), which is presented in this paper as an alternative to LMM.) These models aid in predicting disease risk and drug responses based on genomic data, laying the ground for the realization of precision medicine (Sundin et al., 2018; Park et al., 2022).

To assess the validity of these statistical models, a straightforward approach is to partition the dataset into a training set and a hold-out testing set. The test set is not utilized in model fitting (Nakagawa & Fujita, 2018), therefore serves to test the model’s generalizability. The test error calculated via the application of a trained model over a hold-out set presents certain limitations. This method is not effective when dealing with smaller datasets and lacks the capability to calculate variance with respect to the training set, rendering it unsuitable for algorithm comparisons (Rabinowicz & Rosset, 2022; Dietterich, 1998). As such, the cross-validation (CV) technique, which partitions data into folds and conducts training and testing over each fold, is commonly employed for estimating model generalizability. For instance, in a comparative study of mixed and regularization models, the Lasso and elastic net (ENET) models were selected using 100-fold cross-validation (Zeng et al., 2017).

Using CV to assess the accuracy of models is popular in many fields including genomics-based predictions. However, the estimated ancestral effective population size of humans may be between 12,800 and 14,400 (Henn, Cavalli-Sforza, & Feldman, 2012), leading to the prevalence of relatedness among individuals in any dataset. Even worse, in most cohorts, the relatedness between individuals is uneven. As a result, the validity of cross-validation can be challenged due to the extensive and heterogeneous relatedness among human genes. This relatedness contradicts the identical independent distribution (i.i.d.) assumption that is commonly used for cross-validation and is rarely been studied (Wieczorek, Guerin, & McMahon, 2022). Furthermore, the relatedness leads to a violation of the assumption that the test set in CV can represent unseen data for prediction (Tabe-Bordbar, Emad, Zhao, & Sinha, 2018). However, in the field of biostatistics and genomics, this issue has not been recognized, not to mention its characterization and correction.

To ease the correlation problem in CV, a correction method called CVc is introduced by A. Rabinowicz and S. Rosset (2022). Their research presents a fresh viewpoint on cross-validation and how cross-validation is affected by the correlation structure: the CV is biased if and only if the joint distribution of split folds for training and validating is the same as the joint distribution of the current data with future data for prediction. Based upon this insight, they introduced a bias-corrected measure of CV, referred to as CVc, for validating multiple situations when the cross-validation is biased. However, being a formulation for general CV corrections, the CVc method does not tailor to specific models used in genomic prediction.

In this study, we conducted a comprehensive examination of CV bias across various models, including the Ordinary Least Square (OLS), Generalized Least Squares (GLS) (Hansen, 2007), polygenic method, i.e. LMM with its predictor gBLUP, three regularization methods, i.e. Ridge, Lasso, and ENET (Tibshirani, 1996; Zou & Hastie, 2005; Hastie, Tibshirani, & Wainwright, 2015). Additionally, We also explored two hybrid methods: One combining the GLS with *L*_2_ regularization (GLS+Ridge), the other combining gBLUP with *L*_2_ regularization (gBLUP+Ridge). These hybrid methods estimate the effects by minimizing Mahalanobis distance with a *L*_2_-norm of the coefficients. Specifically, We utilized a genomics dataset created by The 1000 Genomes Project Consortium (Consortium et al., 2015), complemented with simulated phenotypes, for our investigation.

To generate the “gold-standard” dataset that is devoid of bias due to relatedness within the real human genomes, we created a set of synthetic genomes. These synthetic genomes are generated based on the Minor Allele Frequency (MAF) of the real genomes, however, without retaining the linkage disequilibrium (LD) (Slatkin, 2008) present in the real genomes. This approach ensures that the synthetic genomes mirror the alleles distribution of genetic variants found in real genomes, simultaneously, being free from relatedness.

As such, we assume the synthetic genomes can be used to test the real generalization errors. The phenotypes for these synthetic genomes are generated using the same coefficients simulated previously in the main 1,000 Genomes Project data. With the aid of “real” phenotypes on the synthetic genomes, the standard Mean Square Error (MSE) of phenotype predictions on the synthetic genomes using any of the models mentioned above can be calculated, which is viewed as the gold-standard error estimation as the generalizability. The derivation of CV estimation of errors from the gold-standard errors, hence, quantifies the bias of CV. Following this procedure, we compared the CV error with MSE on both the test set and synthetic set as a measurement of the CV bias. Our analysis revealed the biases of CV error compared to gold-standard error for the methods listed in Table 1 in genomic prediction.

**Table 1:**
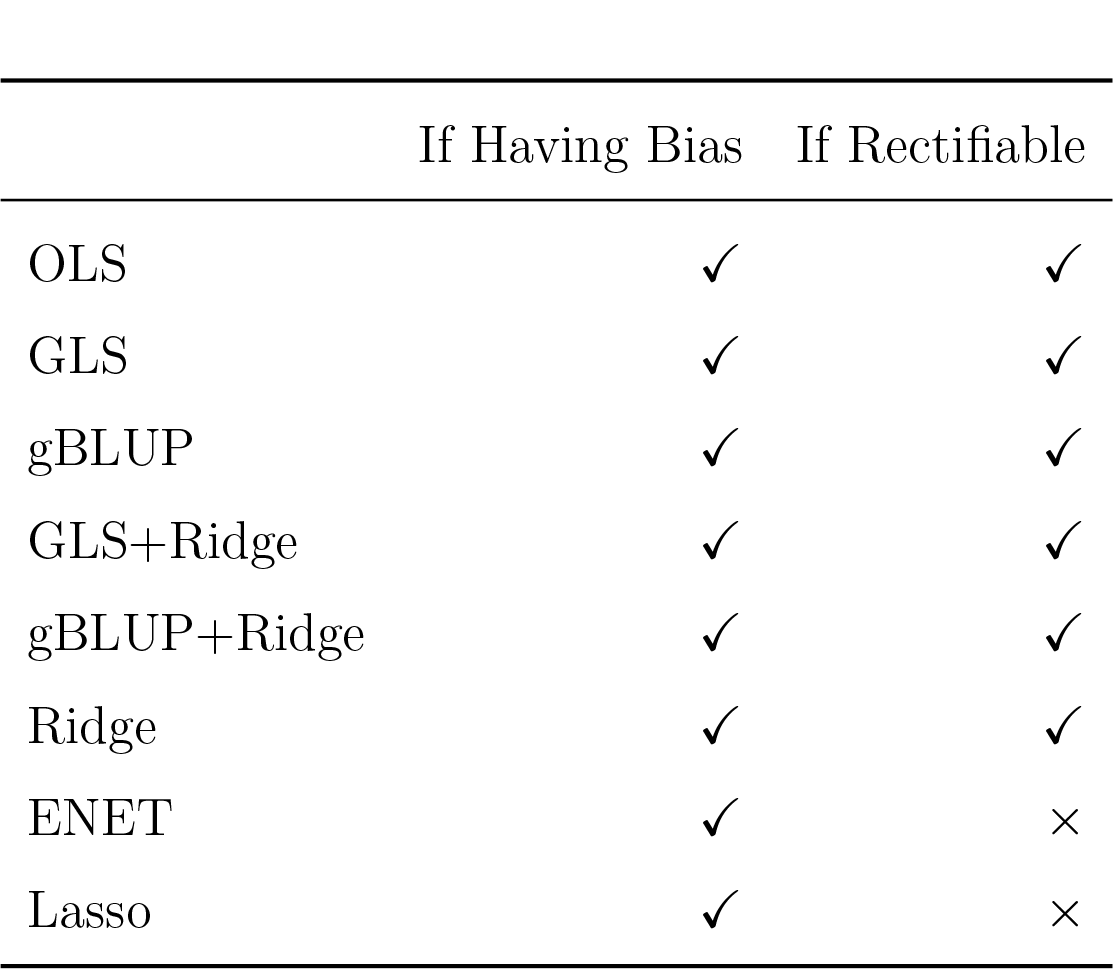
The table shows if the following methods have a bias for CV and if we can correct it.

To rectify these biases, we adapted the aforementioned CVc method by deriving related closed-form formulas tailored to the models under investigation. CVc method calculates the correction by adding the difference of covariance of the predicted dependent variable and the dependent variable in the cross-validation process with the covariance in the testing process. To calculate the covariance, one extracts the projection matrix from the covariance, which means only linear methods with closed-form solutions can be applied to rectify the CV bias. Consequently, we derived the projection matrices for the models in Table 1 by extracting the target variable **y** from their prediction formulas. The derived models were subsequently compared with the original CV model by checking their test errors assessed by the gold standard, our standard estimation of the generalization error. We found that CVc can successfully produce better estimates on the six methods listed in Table 1 with respect to their generalizability assessed on the synthetic set.

## 2 Bias of Cross-validation

In order to demonstrate the bias of cross-validation, we compared the performance of the methods mentioned in Table 1 using simulated phenotype data.

### 2.1 Definition of the Bias

We start by defining the following terms. The K-folds cross-validation error (CV) is defined as follows: given a whole dataset *T* that is randomly partitioned into K folds,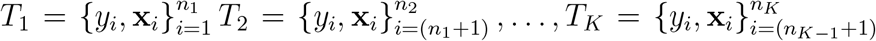, where *n*_*k*_ − *n*_*k*−1_ is the sample size in fold *k* and *n*_*K*_ = *n*. The cross-validation error is then calculated as:

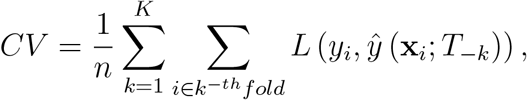

Where ŷ (**x**_*i*_; *T*_−*k*_) is the prediction of *y*_*i*_, constructed by training with *T*_−*K*_ and predicting on **x**_*i*_. Additionally, for each *k* ∈ {1, …, *K*}, the model is trained on the entire data excluding the *k*-th fold, denoted by *T*_−*k*_ =⋃_*j*/=*k*_ *T*_*j*_ = {**y**_−*k*_, *X*_−*k*_}. The prediction error of the trained model is then measured on the holdout fold, *T*_*k*_ = {**y**_*k*_, *X*_*k*_}, with respect to some loss function *L*(·, ·) : R × R → R. Especially, when *K* = *n*, the method is called leave one out (LOO) cross-validation.

The CV estimator is used to estimate the generalization error, which reflects the model’s ability to generalize. Let the training set be denoted as 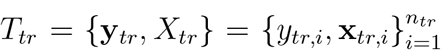 and the test set be denoted as 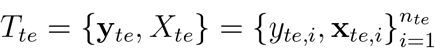 The generalization error is defined as:

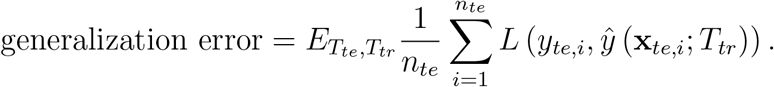

The generalization error is estimated by applying the trained models to both the test set and the synthetic set. In this study, MSE for the trained model applied to the synthetic set is considered as the gold standard.

The bias of cross-validation is defined as the difference between the generalization error and the cross-validation error (Rabinowicz & Rosset, 2022). Mathematically, it is expressed as:

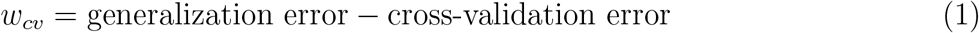

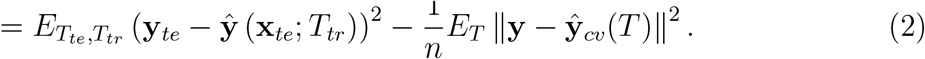

### 2.2 Data Simulation

We employed simulation methods on a real genomics dataset to emulate authentic data closely. Specifically, we leveraged the dataset from the 1000 Genomes Project (Consortium et al., 2015), encompassing whole genome sequencing across diverse populations. This dataset provides 2,504 samples which are sampled from 26 populations from five continental groups. In order to reduce the influence of rare variants that weaken the model’s ability, we first filtered out Single Nucleotide Polymorphisms (SNPs) with a Minor Allele Frequency (MAF) smaller than 0.05. We then randomly selected a total of *p* = 10, 000 SNPs in the first chromosome. Moreover, all the SNPs we chose are from nearby genes and therefore are considered cis-SNPs (Zeng et al., 2017). The phenotype was simulated using the following formula:

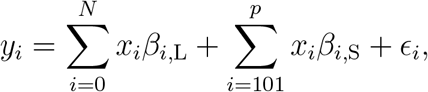

where *N* is the number of large effects 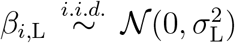 are the large effects 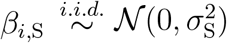 are the small effects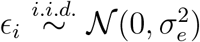 To test the CV bias in different scenarios,we Set *N* = 100 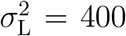 and 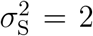, which means We sampled 100 SNPs with large effects and the remaining SNPs are small effects therefore to simulate the situation that all SNPs in **X** has strong effects.

The genotypes that we sampled are from the same chromosome and therefore have nonnegotiable LD. To showcase the model’s efficacy on ground truth that is independent of the current data and free from LD as shown in Figure 1a, we generated synthetic genomes. These synthetic genomes are devoid of the LD present in real data, serving as a gold standard for evaluation. This synthetic data comprises 10,000 SNPs and *n*_*te*_ samples. Each SNP was simulated independently, based on its MAF derived from real genomic data. Let *p*_*i*_ denote the MAP at loci *i*. We set the probability of SNP at loci *i* being minor variants as *p*. Therefore the probability of homozygote of both minor variants is *p*^2^, the probability of heterozygote is 2*p*(1 − *p*), and the probability of homozygotes of the reference variants is (1 − *p*)^2^. Subsequently, phenotypes were generated using the effects we had previously determined. Since every SNP in the synthetic set is generated independently, the frequency *p*_*AB*_ = *p*_*A*_*p*_*B*_, in other words, the LD for any two different SNPs in the synthetic set is zero as shown in Figure 1b. In the synthetic set, each SNP is generated independently. As a result, the joint frequency of two alleles, *p*_*AB*_, is simply the product of their individual frequencies: *p*_*AB*_ = *p*_*A*_*p*_*B*_. Consequently, there is no linkage disequilibrium (LD) between any pair of distinct SNPs in the synthetic set, as depicted in Figure 1b.

**Figure 1:**
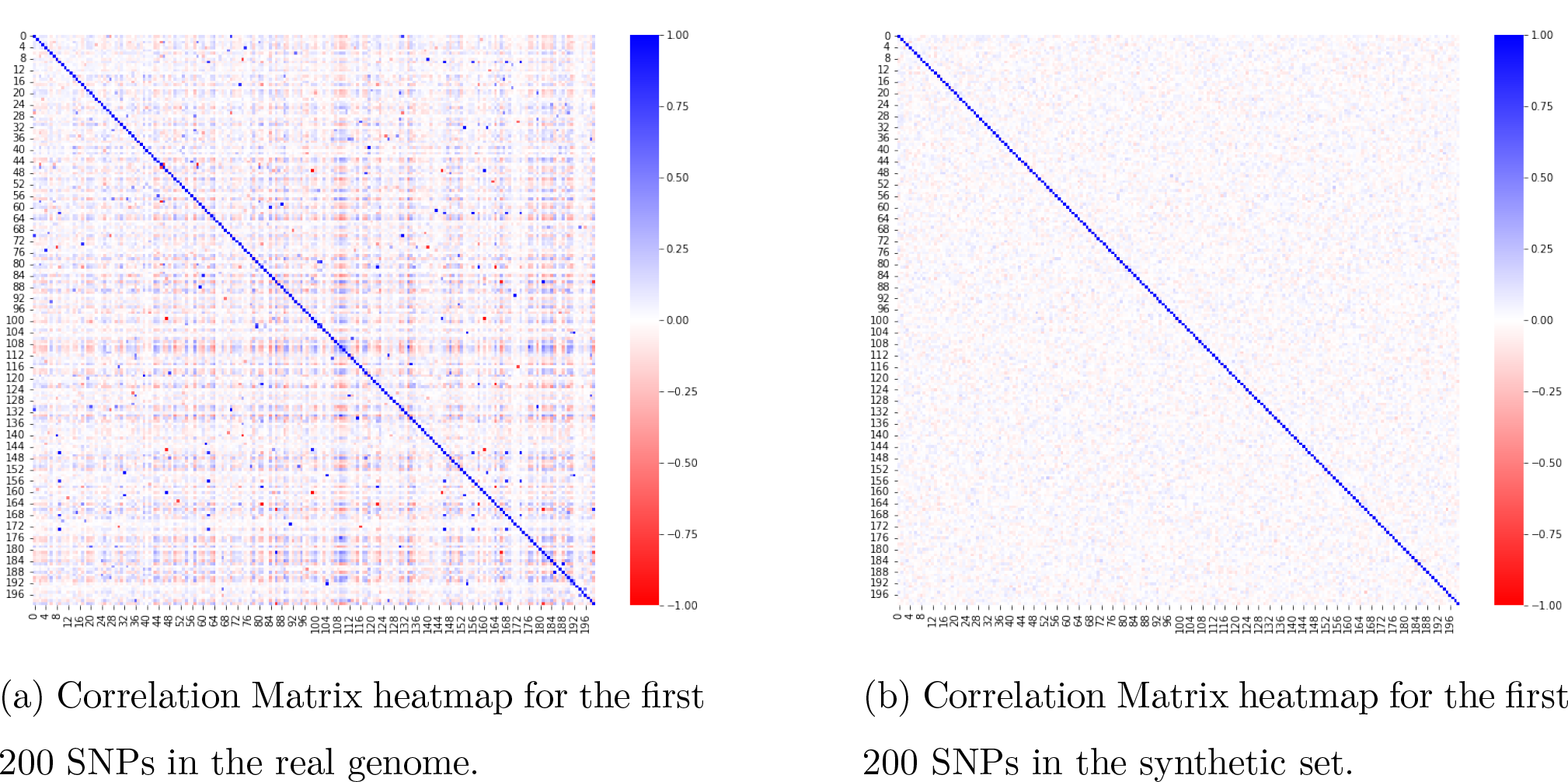
The correlation matrix of first 200 SNPs in the selected real genome and synthetic set. The synthetic set eliminates the correlation.

The synthetic genomes mirror the effects seen in real genomic data while being devoid of LD. This synthetic set serves as a tool for evaluating the model’s generalization ability. Given the absence of SNP correlations in this set, issues related to data leakage during crossvalidation fold splitting are eliminated. As such, a model’s performance on this synthetic set can be considered the gold standard.

In our simulation study, we carried out 100 replications. For every replicate, the genomic effects and residual effects are independently sampled and thus every replicate is identical and independent of each other.

### 2.3 The Bias of Cross-validation

As depicted in Figure 2, the MSE for the test set is similarly distributed as the crossvalidation error, whereas the gold standard is significantly higher than the CV. Figure 2b illustrates the distribution of the difference between the MSE of the test set and the CV error, and the difference between the gold standard and the cross-validation error, highlighting the lack of generalization ability of these methods except GLS. Notably, while gBLUP+Ridge exhibits the lowest CV error, Lasso achieves the minimum test error on the synthetic set.

**Figure 2:**
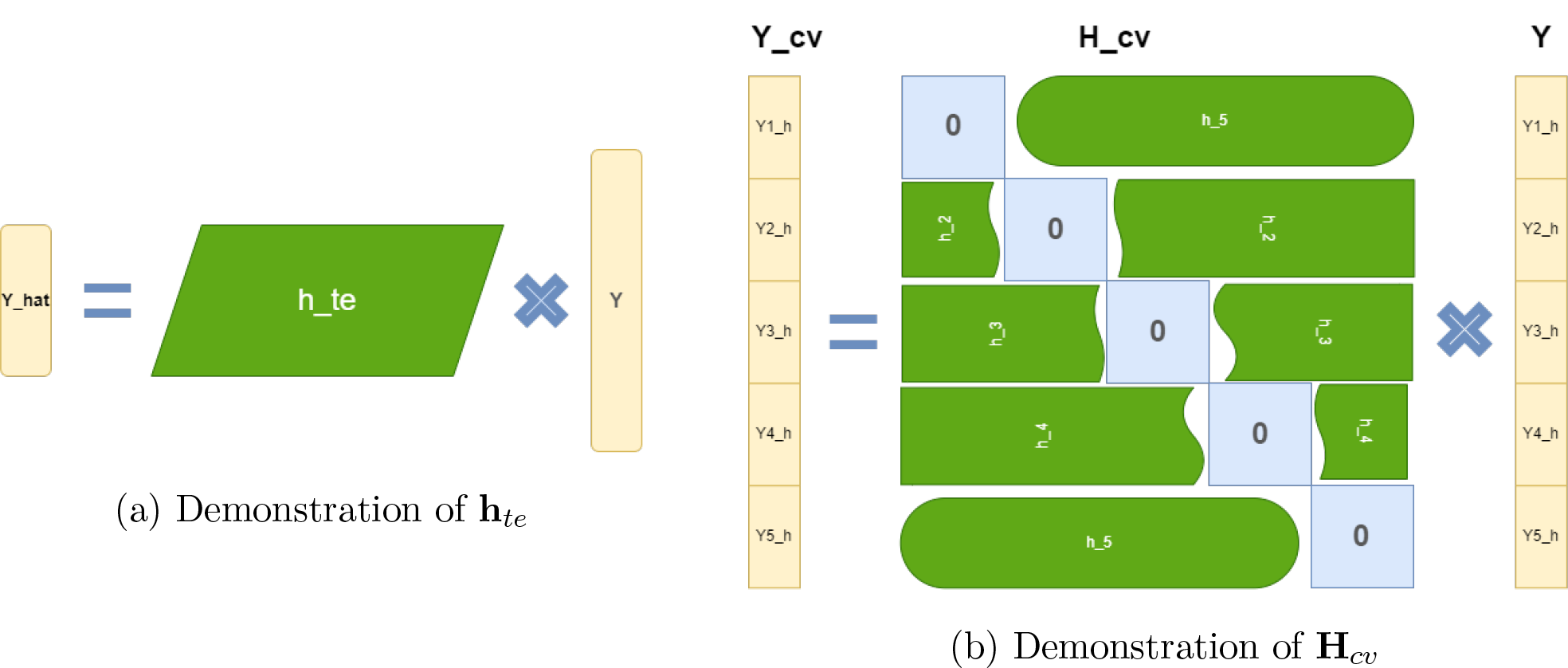
Demonstration of Projection Matrix: (a) *h*_*te*_ is a linear transformation matrix that transforms the known target to prediction. (b) The graph illustrates the 5-fold CrossValidation. *H*_*cv*_ has all diagonal matrices as zero.

**Figure 3:**
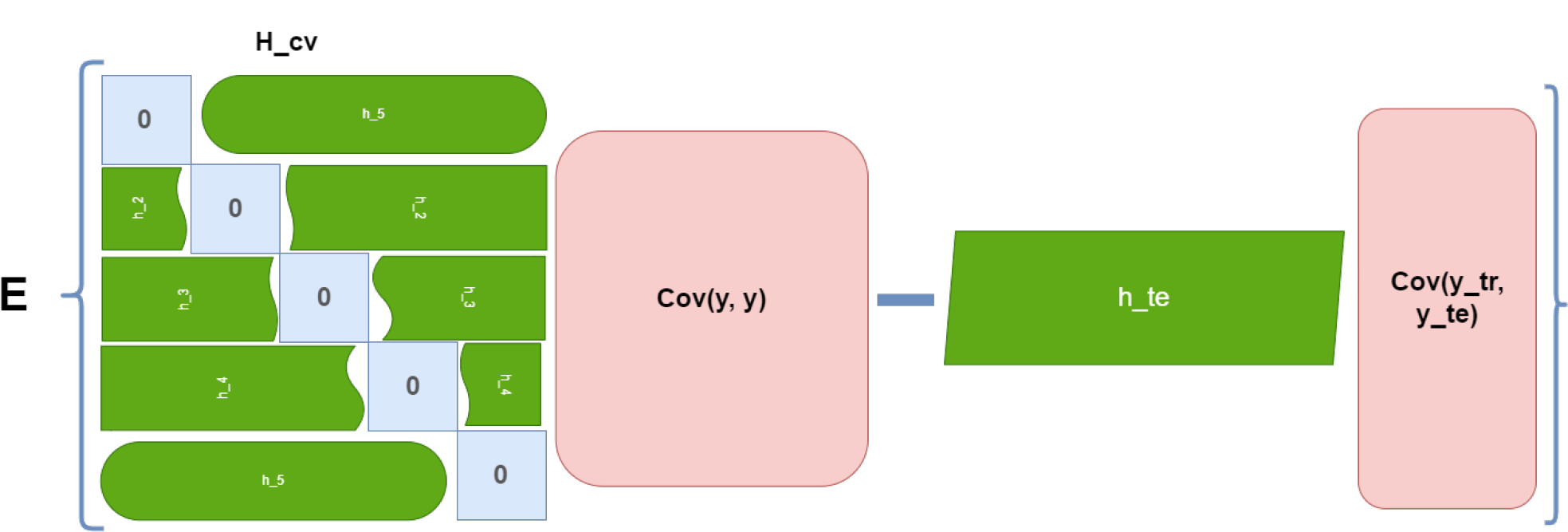
Demonstration of calculation of *w*_*cv*_

## 3 Correction Methods

To correct the bias, we employed a technique known as **cross-validation correction (CVc)** (Rabinowicz & Rosset, 2022). This method provides an unbiased estimate of the generalization error when cross-validation is given. Before we delve into the correction process, it is essential to define linearity in a machine learning model and introduce a projection vector that enables the ŷ to be expressed as a linear combination of **y**, since the method can be only applied to the models that are considered as linear.

### Definition 3.1.

A machine learning model is termed “linear” when the predicted dependent variable of a test sample, ŷ (***x***_*te*_) ∈ R, can be expressed as a linear combination of the dependent variable 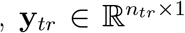 in the training process, where ***x***_*te*_ ∈ ⊮ × | is vector of predictors for a testing sample. Mathematically, let’s consider a vector **h**_*te*_ ∈ R^1×*n*^. A model is defined as linear if its predictions can be represented in the form

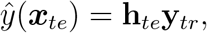

where **h**_*te*_ is constructed by ***x***_*te*_, training data **X**_*tr*_ and 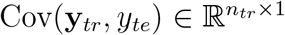 Such vector **h**_*te*_ is called a projection matrix. When the predicted target 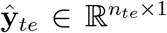 is a vector of multiple predicted dependent variables using the trained model, the projection vector can be stacked to become a projection matrix.

This definition implies that for a test sample, its prediction is a linear combination of all training samples. When there are multiple samples to be predicted, the Projection Vector becomes a projection matrix 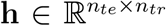, where *n*_*te*_ is the number of test samples and *n*_*tr*_ is the number of training samples. If a model prediction can be represented as such a form of linear combination, it is called **Linear**.

### Definition 3.2.

**H**_*cv*_ ∈ R^*n*×*n*^ is a projection matrix shown in Figure 2b for **y** if **ŷ**_*cv*_ ∈ R^*n*×1^ is a linear combination of **y** ∈ R^*n*×1^ and

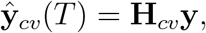

where **H**_*cv*_ ∈ R^*n*×*n*^ does not contain **y** and is constructed as follows:

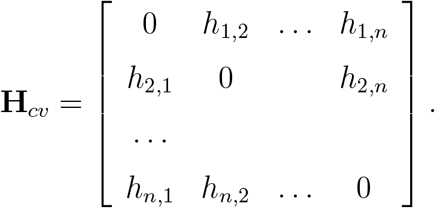

In this definition, **H**_*cv*_ is a matrix with all diagonal elements set to zero. It maps **y** to the prediction ŷ_*cv*_ in LOO cross-validation. This implies that for each sample, its prediction is a linear combination of all other samples, excluding the sample itself. For instance, when the diagonal element zero is removed, the *k*-th row is a projection vector **h**_*k*_ ∈ R^1×(*n*−1)^ which is defined in Definition 3.1 and constructed from **X**_−*k*_ and covariance Cov(**y**_−*k*_, *y*_*k*_) ∈ R^(*n*−1)×1^. The projection vector linearly combines all the sample responses other than the *k*^th^ sample *y*_*k*_.

In the case of k-fold cross-validation, **H**_*cv*_ is constructed similarly to the LOO method, except that the diagonal positions contain block matrices of zero instead of individual zero entries. The projection matrix of the *k*^th^ fold, denoted as **h**_*k*_ ∈ R^*f* ×(*n*−*f*)^, is constructed in a similar to the LOO method, where *f* = *n/K* represents the number of samples for each fold.

According to the paper (Rabinowicz & Rosset, 2022), the CV correction denoted as *w*_*cv*_, is given by:

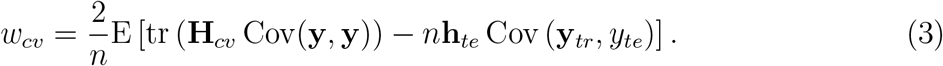

In situations where we have more than one testing data, the test projection matrix is considered as a matrix 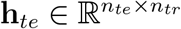 . Thus, the calculation of *w*_*cv*_ becomes:

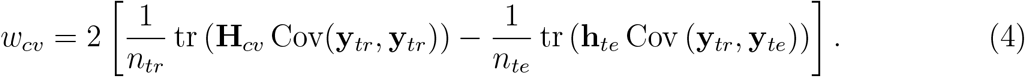

This correction for CV is intuitive because it calculates the differences between the covariance of training and testing data and the training data itself.

### 3.1 Construct the Projection Matrix

To apply the correction method, we need to determine the linear combinator, **H**_*cv*_, for different machine learning methods.

Let’s start with the simplest method, linear regression. Considering a regression model:

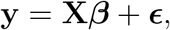

where **X** is the design matrix of the marker effects and the relatedness matrix 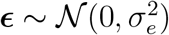 is the vector of residuals.

The estimation of the parameter 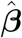 for OLS is given by 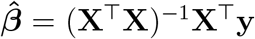 Therefore the prediction using OLS for the test set is

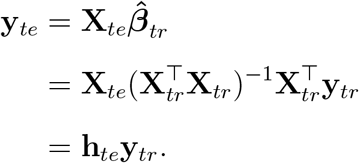

Therefore the construction for the project matrix for testing is

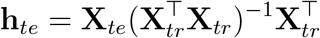

Similarly, the construction of **H**_*cv*_ for the *k*^th^ fold’s projection matrix **h**_*k*_ is as follows:

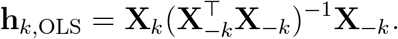

Regularization methods like Ridge can also be formulated in this way. Ridge Regression adds an *l*^2^ norm to the sum of squares of residuals to form a loss function. By minimizing this function, we get the estimate for 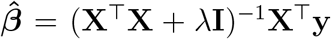 Therefore the prediction using Ridge for the test set is

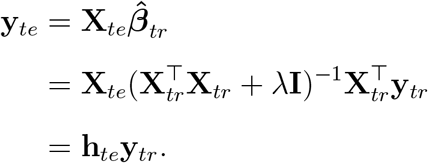

Therefore the construction of the projection matrix for testing is

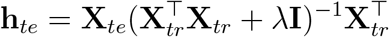

Similarly, the *k*^th^ fold projection matrix is:

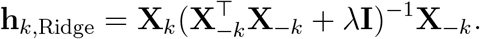

The construction of the projection matrix for the above models does not involve the covariance of the response variable because these models do not include correlation in the model and assume the i.i.d assumption for the samples. However, in the models, we will discuss next, the variance structure is included in the model.

Let us consider an LMM, represented as:

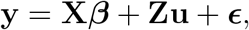

with 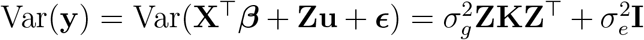 where **X** is the design matrix of the marker effects and the relatedness 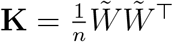, where 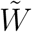 is standardized genomic *n* × *s*_*c*_ matrix *W* where *s*_*c*_ is the number of SNPs that is used to calculate the relatedness matrix. *W* is a genomic matrix with *W*_*ij*_ ∈ {0, 1, 2} indicating the number of variants in loci *j* for the *i*^*th*^ individual. And 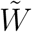 is calculated as

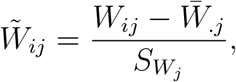

where 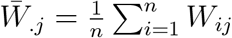 the sample mean and 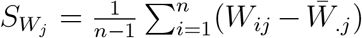 is the sample variance.

Given the variance components 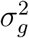 and 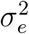 estimated either by Restricted Maximum Likelihood (REML) or Maximum Likelihood Estimate (MLE), one of the solutions for a heteroscedasticity linear model is GLS 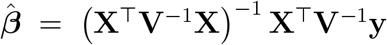prediction using Ridge for the test set is

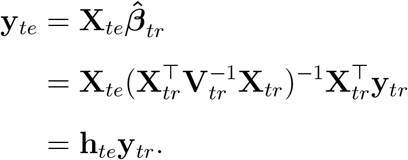

Therefore the construction of the projection matrix for testing is

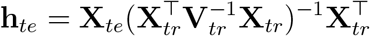

Thus the *k*^th^ fold projection matrix is

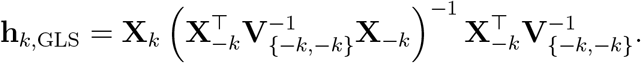

The estimator for the random variable is called the genomics best linear unbiased estimator (gBLUP). To get gBLUP for the test set, let us consider the joint normal distribution of **u**_*te*_ and **y**_*tr*_ can be written as

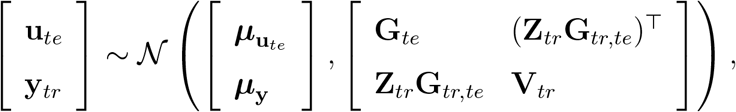

Using the property of joint normal distribution, we can compute the conditional expectation of the random vector given the predictors **y**, and therefore

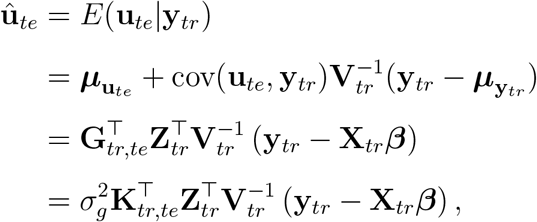

where 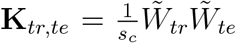 is *n*_*tr*_ × *n*_*te*_ matrix. We can replace the ***β*** in the gBLUP with

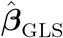allowing us to extract the **y** from gBLUP

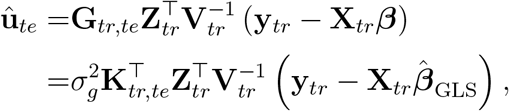

Since the estimator will be fixed effects estimated by GLS added up with gBLUP, i.e.,

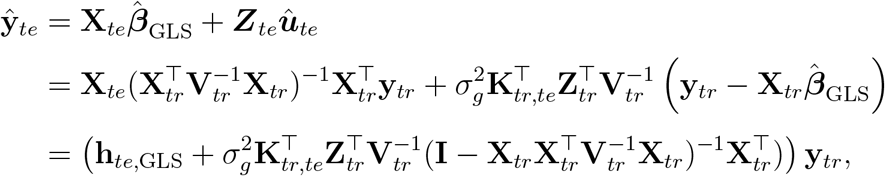

the estimator for the fixed effects and random effects in an LMM can be computed using the GLS method and the gBLUP method, respectively. The combined projection matrix or predict matrix for the LMM is then given by:

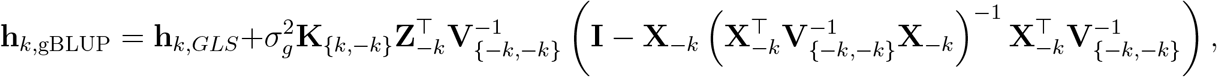

where **K**_{*k*,−*k*}_ is the relatedness matrix between the *k*^th^ folds and the data without *k*^th^ folds.

Similarly, L2 regularization can also be added when fitting the LMM. We estimate the *β* by minimizing the Mahalanobis distance with a *l*^2^-norm of the *β*, i.e.

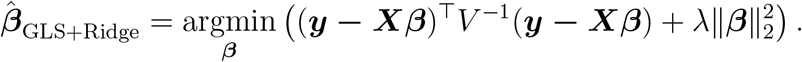

Thus, the estimate for *β* is 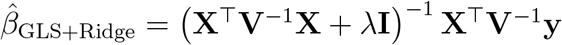 Thus, the prediction for the test set using GLS+Ridge is

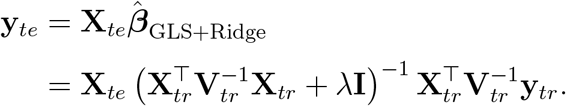

Then, we get the projection matrix for GLS+Ridge is

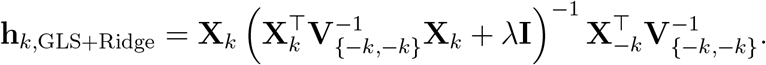

Consequently, the projection matrix for gBLUP+Ridge, i.e.,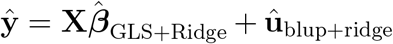 can be calculated by plugging in 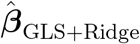 into û_gBLUP+Ridge_:

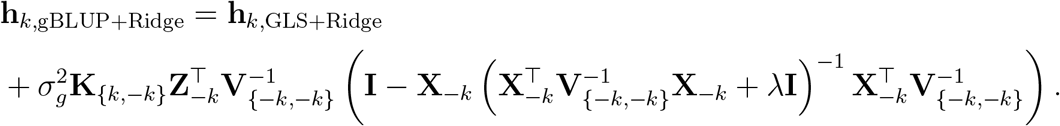

The projection matrices for the methods that are shown in Table 1 are

- **OLS**: Prediction Matrix is

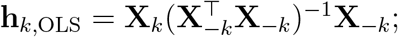

- **GLS**: Prediction Matrix is

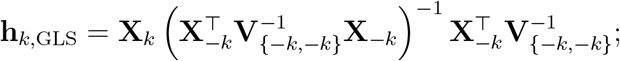

- **gBLUP**: Prediction Matrix is

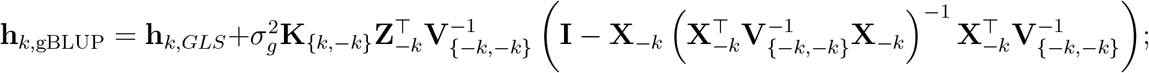

- **Ridge**: Prediction Matrix is

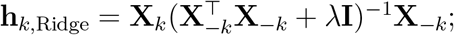

- **GLS+Ridge**: Prediction Matrix is

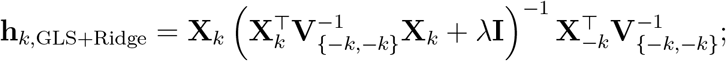

- **gBLUP+Ridge**: Prediction Matrix is

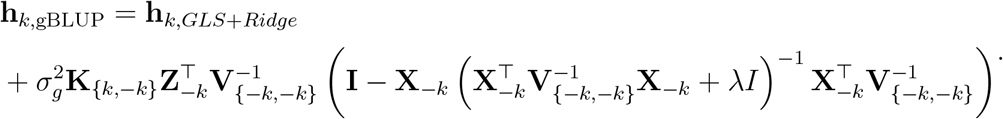

## 4 Assessing the Bias Corrections by Synthetic Data

We applied the CVc to the simulation data. Given that there are no analytic solutions for the regularization methods ENET and Lasso, we compared the correction for the other six methods on both the test set and the generated data. We applied CVc to it as well to examine how the method performs in a less biased situation.

The correction distribution in Figure 4a demonstrates a significant correction effect for the synthetic set especially for the gBLUP-based model: both gBLUP and gBLUP+Ridge got more than 60% bias corrected as shown in the Table 2 .

**Table 2:** The CV error, MSE, CVc, and percentage of the correction on both test set and gold standard for the 8 methods (Lasso and ENET do not have correction).

|  | OLS |  |  | GLS |  |  | gBLUP |  |  | GLS+Ridge |  |  | gBLUP+Ridge |  |  | Ridge |  |  | ENET |  |  | LASSO |  |
| --- | --- | --- | --- | --- | --- | --- | --- | --- | --- | --- | --- | --- | --- | --- | --- | --- | --- | --- | --- | --- | --- | --- | --- |
|  | mean | std |  | mean | std |  | mean | std |  | mean | std |  | mean | std |  | mean | std |  | mean | std |  | mean | std |
| CV | 0.86 | 0.20 |  | 1.09 | 0.22 |  | 0.74 | 0.16 |  | 0.98 | 0.19 |  | 0.72 | 0.15 |  | 0.92 | 0.17 |  | 1.36 | 0.36 |  | 0.84 | 0.17 |
| MSE<sub>test</sub> | 0.86 | 0.19 |  | 1.09 | 0.23 |  | 0.74 | 0.17 |  | 0.98 | 0.19 |  | 0.70 | 0.15 |  | 0.92 | 0.19 |  | 1.36 | 0.38 |  | 0.84 | 0.20 |
| MSE<sub>synthetic</sub> | 2.03 | 1.07 |  | 2.56 | 2.62 |  | 2.05 | 1.08 |  | 2.33 | 1.41 |  | 1.95 | 1.24 |  | 2.36 | 1.69 |  | 2.39 | 2.08 |  | 1.36 | 0.89 |
| CV<sub>Ctest</sub> | 0.86 | 0.20 |  | 1.10 | 0.22 |  | 0.72 | 0.16 |  | 0.99 | 0.18 |  | 0.70 | 0.15 |  | 0.93 | 0.17 |  |  |  |  |  |  |
| CV<sub>Csynthetic</sub> | 1.28 | 0.24 |  | 1.33 | 0.25 |  | 1.53 | 0.26 |  | 1.18 | 0.20 |  | 1.52 | 0.24 |  | 1.39 | 0.22 |  |  |  |  |  |  |
| Correction<sub>synthetic</sub> (%) | 35.39 | 5.52 |  | 16.30 | 2.41 |  | 60.24 | 10.33 |  | 15.04 | 1.59 |  | 64.93 | 9.80 |  | 32.31 | 5.13 |  |  |  |  |  |  |

**Figure 4:**
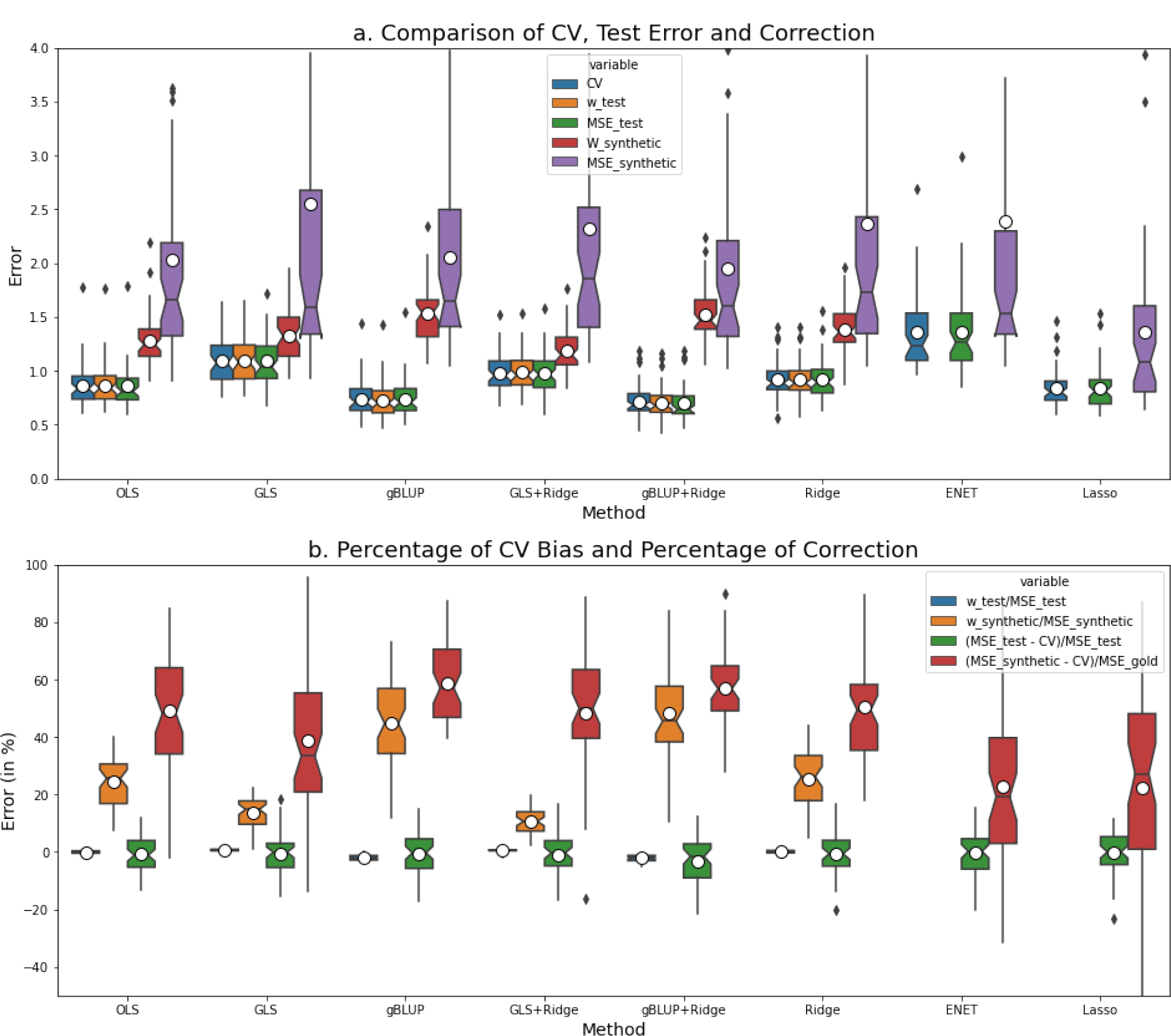
Bias and Correction: a: The distribution for CV, MSE test, MSE gold standard, and CVc. b: The blue and yellow box plots represent the correction percentages for the test set and synthetic set, respectively. Meanwhile, the green and orange boxplots depict the biases for the test set and synthetic set, respectively.

Furthermore, Figure 4b displays the distribution of the ratio correction with the MSE for both the test set and gold standard respectively. It illustrates the averaged zero correction for the test set and positive correction for the synthetic set that is devoid of LD. Interestingly the GLS-based algorithms have high variances of CV and MSE for both the test set and generated data, both with Ridge regularization and without Ridge regularization with Ridge. This phenomenon could possibly explain the non-significance of the CV bias, as the model already has a large variance that can be shown by CV.

Among these methods, while the CVc method cannot significantly adjust the GLS-based model, the gBLUP-based model gives the best performance in prediction and gBLUP + Ridge further reduces the spreading on predicting the generated data and gives the lowest MSE on the test set and the gold standard.

## 5 Discussion

In this paper, we delved into the potential underestimation of the true error by the CV estimator due to the inherent correlation of genomic data among individuals. We evaluated the generalization capabilities of several prevalent methods for predicting gene expression, including OLS, GLS, gBLUP, Ridge, Lasso, and ENET. Additionally, we examined two hybrid methods: GLS combined with Ridge regularization and gBLUP combined with Ridge regularization. Our findings indicate that while cross-validation offers a reliable estimate of the test error on a designated test set, it tends to significantly underestimate the prediction error for datasets that is independent of the training set.

To illustrate this underestimation, we employed a synthetic set, generated independently for each SNP. This synthetic set, devoid of Linkage Disequilibrium (LD), serves as a ground truth for the methods under consideration.

To address this issue, we adapted a correction method to the CV, known as CVc, which utilizes the covariance of the training set and test set. This correction significantly improved the accuracy of predicting the test error for all mentioned methods except Lasso, and ENET based on the projection matrix we derived. The correction method performs the best when correcting cross-validation for gBLUP-based models.

However, there are certain limitations in our study. Firstly, using the model performance on data generated with the MAF as the gold standard for testing the model may not be entirely representative. The synthetic set is designed to eliminate the influence of LD which causes the data leakage therefore underestimation of the generalization error. Given the presence of LD in the actual human genome, the model performance on the synthetic set might not fully emulate the real human genome.

Secondly, this study did not delve into the selection of the regularization hyper-parameter. We adhered to a fixed hyper-parameter setting, even though variations in this parameter could affect the model correction.

Finally, the CVc approach necessitates the model to be linear with a closed-form solution for constructing the projection matrix. This poses a limitation because non-linear methods, such as decision tree-based techniques, are not amenable to rectification. Furthermore, linear models using Bayesian approaches are not rectifiable due to the absence of closed-form solutions.

Consequently, the widely adopted Bayesian Sparse Linear Mixed Model (BSLMM) (Zhou et al., 2013) in the genomics field was not incorporated into our analysis. This omission arises from the lack of analytical correction solutions in Bayesian methods. Implementing cross-validation for Bayesian models also poses challenges, as each fold requires running a new MCMC chain for parameter estimation causing computational intensive.

Going forward, to adapt CVc in BSLMM, we intend to leverage an equivalence between Bayesian models and Frequentist Mixed Models. This would facilitate the transformation of a Bayesian model into a mixed model, enabling us to correct cross-validation bias.

In conclusion, this paper underscores the potential pitfalls of relying solely on crossvalidation or hold-out prediction errors, especially when correlations exist among samples. This issue is particularly important to consider when validating a model, especially in scenarios in genomic prediction due to the existence of LD. In such contexts, The CVc method is effective as it uses the covariance on training data, subtracting the small correlation on the test set to correct the CV.

## SUPPLEMENTARY MATERIAL

**Code:** The source code for the Cross-Validation correction and simulation script can be found on https://github.com/theLongLab/CVc_in_bioinformatics.

